# Hard decisions shape the neural coding of preferences

**DOI:** 10.1101/298406

**Authors:** Katharina Voigt, Carsten Murawski, Sebastian Speer, Stefan Bode

**Affiliations:** Melbourne School of Psychological Sciences, The University of Melbourne, Victoria 3010, Australia; Department of Finance, The University of Melbourne, Victoria 3010, Australia; Rotterdam School of Management, Erasmus University, 3000 DR Rotterdam, The Netherlands; Department of Psychology, University of Cologne, 50969 Cologne, Germany

**Author notes:** Corresponding author: Katharina Voigt The University of Melbourne Redmond-Barry Building Parkville, VIC 3010, Australia. **Author Contributions:** KV, CM, SB, and SS contributed to the study design. KV and SS performed testing and data collection. KV conducted data analyses. KV and SB wrote the paper and all authors approved the final version of the manuscript for submission.

**Keywords:** Preference formation, choice-induced preference change, decision-making, dorsolateral prefrontal cortex, eye movements

## Abstract

Hard decisions between equally valued alternatives can result in preference changes, meaning that subsequent valuations for chosen items increase and decrease for rejected items. Previous research suggests that this phenomenon is a consequence of cognitive dissonance reduction after the decision, induced by the mismatch between initial preferences and decision outcomes. In contrast, this functional magnetic resonance imaging and eye-tracking study tested whether preferences are already updated online while making decisions. Preference changes could be predicted from activity in left dorsolateral prefrontal cortex and precuneus during decision-making. Furthermore, fixation durations predicted both choice outcomes and subsequent preference changes. These preference adjustments became behaviourally relevant at re-evaluation, but only for choices that were remembered and were associated with hippocampus activity. Our findings refute classical explanations of post-choice dissonance reduction and instead suggest that preferences evolve dynamically as decisions arise, potentially as a mechanism to prevent stalemate situations in underdetermined decision scenarios.

## Introduction

Traditional neurocognitive models of value-based choice viewed decision-making as a serial process in which *stable* preferences are the basis of subsequent choices (Dolan and Dayan, 2013). However, there are decision scenarios in which choice options appear equally valuable to the decision-maker, and therefore existing preferences are not sufficient to rank alternatives. In Jean Buridan’s philosophical parable, a hungry donkey is placed between two bales of hay. As both choice options appear equally appealing, the donkey is unable to decide and eventually starves to death. This parable tells us that there are hard decisions, in which existing preferences are not sufficient to identify a preferred option. Instead, preferences might need to be re-constructed dynamically online as hard decisions arise if we do not want to end up like Burdian’s starving donkey.

Indeed, substantial evidence suggests that our preferences are not rigid, but evolve dynamically and are dependent on the decision context (Lichtenstein and Slovic, 2006). One highly debated question in decision science is whether the act of choosing among equally valued alternatives (henceforth *hard decisions*) itself shapes preferences. Consider a person deciding between two flavours of ice-cream but initially being indifferent between them. The choice-induced preference change effect refers to the phenomenon that after having made a choice, the chosen option is preferred more, while the alternative is preferred less (Izuma and Murayama, 2013). Prominent explanations of this effect are based on Festinger’s (1957) theory of cognitive dissonance, which proposes that discrepancies between actions (i.e., rejecting a liked ice cream flavour) and preferences causes psychological discomfort.

Preferences are then adjusted after a hard decision has been made to reduce the dissonance between initial preference and the decision outcome (Harmon-Jones, Harmon-Jones, and Levy, 2015 for review). This explanation is in line with neuroimaging studies, which suggested that at the time of re-evaluation, after dissonance between preferences and choices is detected by the anterior cingulate cortex (ACC) (Kitayama, Chua, Tompson, and Han, 2013; van Veen, Krug, Schooler, and Carter, 2009) the dorsolateral prefrontal cortex (dlPFC) triggers changes in the underlying neural representation of value (Izuma et al., 2010; Izuma et al., 2015; Mengarelli, Spoglianti, Avenanti, and di Pellegrino, 2013) in the ventromedial prefrontal cortex (vmPFC) or ventral striatum (vStr) (Chammat et al., 2017; Izuma et al., 2010).

An alternative possibility is that preferences are adjusted much earlier, that is, at the time when a hard decision is made, when the value differential of the options is not sufficient to choose among them. As such, preference adjustments might constitute a necessary adaptive (online) mechanism to deal with hard choices, as opposed to a post-decisional process for eliminating cognitive dissonance (Izuma et al., 2010; Izuma et al., 2015). This new hypothesis, however, remains largely untested as existing functional neuroimaging studies (focusing on methodological improvements of the original paradigm; Chen and Risen, 2010) focused entirely on the neural mechanisms of preference change during re-evaluation (Chammat et al., 2017; Izuma et al., 2010).

Based on our hypothesis, we predicted that preference changes will already occur, and are reflected in neural activity, while individuals are making hard choices. We hypothesised that the blood-oxygen-level dependent (BOLD) signal in the dlPFC would predict subsequent preference changes during the deliberation process. Other recent studies suggested that post-decisional preference changes only occur when choices are explicitly remembered (Salti, Karoui, Mailet, and Naccache, 2014), which was associated with left hippocampus activity (Chammat et al., 2017). We therefore also predicted similar effects for preference changes during decision-making. Additionally, we hypothesised that fixation durations play a significant role in solving hard decisions. Support for this conjecture stems from studies demonstrating that visual fixations causally relate to value-based choices: options that are looked at longer are more likely to be chosen (e.g., Krajbich, Armel, and Rangel, 2010) and experimental manipulations of exposure duration bias preferences towards the longer presented option (e.g., Shimojo, Simion, Shimojo, and Scheier, 2003). These findings indicate that future values of choice options might be reconstructed by information gathered ‘in the moment’ via fixations.

In order to test whether preferences change during hard decisions, we used a variant of the *incentive-compatible free choice paradigm* (Voigt, Murawski, and Bode, 2017) while functional magnetic resonance imaging (fMRI) was acquired and eye movements were recorded (*Material and Methods*). Participants (*N* = 22; 13 females; age 18 to 37 years; *M* = 23.57, *SD* = 4.93) first indicated their willingness-to-pay (WTP; between $0-$4) for familiar, well-known and liked supermarket snack food items (valuation phase 1; Fig.1A) outside the scanner, and their bids were used later in an auction to determine which item they obtained for consumption after the experiment. The WTP procedure allowed us to combine similarly valued items (on average *n* = 38.88; *SD* = 4.03) to choice pairs (“hard choices”) for which preference changes were expected (e.g., Izuma et al., 2010; Voigt et al., 2017), while using the remaining items to construct (on average *n* = 19.62; *SD* = 1.20) control pairs (“easy choices”) (Supplemental Material). In the scanner, participants made binary choices between these items from half of the pairs (decision phase 1; Fig.1B). Then, they re-evaluated the items again (valuation phase 2, in the scanner) using an identical WTP procedure. Finally, participants engaged in a second decision phase (in the scanner) in which the other half of choice pairs was presented. This served as a control sequence, assessing (and controlling for) changes in valuation attributable to regression-to-the-mean (Chen and Risen, 2010) (*Material and Methods*). The *spread of alternatives*, calculated as the *difference* between the experimental sequence (valuation-choice-valuation; VCV) and the control sequence (valuation-valuation-choice; VVC) in the *WTP change* from the first to the second valuation phase constituted the choice-induced preference change effect (Chammat et al., 2017). After the experiment, participants completed a choice memory task in which they were shown all items again and had to indicate whether they had chosen or rejected the item, and whether their answer was based on *remembering* this item, or whether they *guessed* their answer based on their current preferences. In addition, eye movements were measured for the majority of our participants (*Material and Methods*) in order to extract fixation durations in the lead-up to a decision in the crucial decision phase 1.

**Figure 1.**
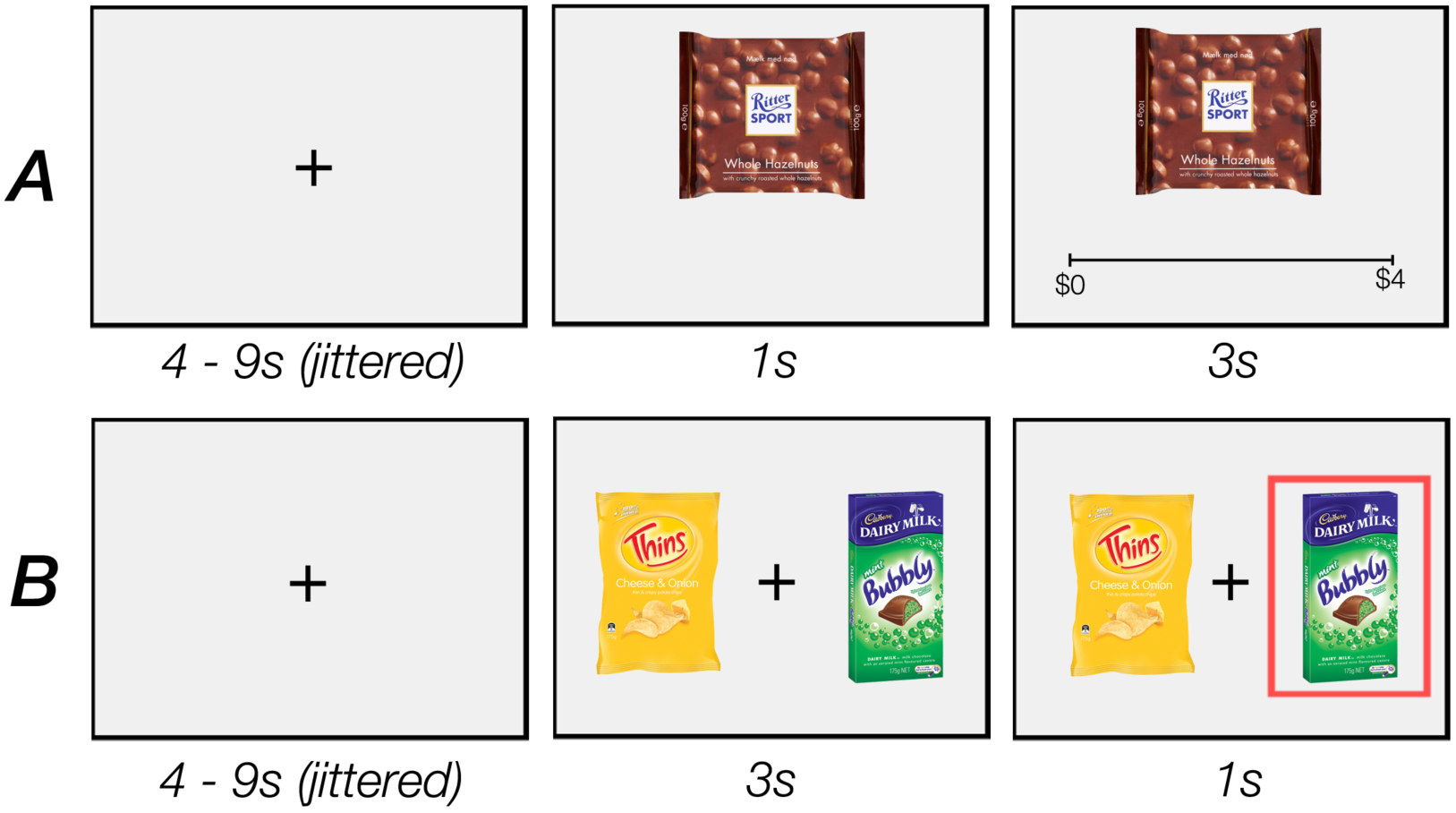
The incentivised free-choice task consisted of four consecutive phases: valuation phase 1, decision phase 1, valuation phase 2, decision phase 2. This task was followed by a choice memory task (not shown). (A) Example trial of the valuation phase. (B) Example trial of the decision phase.

## Results

### Behavioural Results

First, we established the degree of preference changes after individuals made hard choices (i.e., at the time of re-evaluation) and their relation to memory of previous choice outcomes. We found that individuals took significantly longer to decide between equally valued alternatives (“hard decisions”) (*M* = 1.65s, *SD* = 0.27s) than between items that were distinct in their values (“easy choices”) (*M* = 1.41s, *SD* = 0.24s) (*t*(21) = −6.25, *p* <.001).Participants correctly remembered 32.63% (*SD* = 9.41%) and correctly guessed 25.81% (*SD* = 2.40%) of the choice outcomes of their hard choices (difference *n.s., t*(21) = 1.71, *p* >.10). The interaction between experimental condition (i.e., VCV, VVC) and choice memory is considered diagnostic of a memory-dependent choice-induced preference changes (Chammat et al., 2017; Salti et al., 2014) such that only correctly remembered items should show preference changes, but not guessed or not remembered items. In a linear mixed effect (LME) model analyses (Supplemental Material), this interaction showed the hypothesised post-choice spread in valuations for items that were correctly remembered (Condition x Memory interaction effect, *β* = 0.38, *SE* = 0.16, *p* <.02) but not for correctly guessed items (Condition x Guessing interaction effect, *β* = −0.06, *SE* = 0.09, *p* = .53 (Fig. S2). Replicating our previous behavioural results (Voigt et al., 2017), this interaction effect was bidirectional, i.e., WTP increased for correctly remembered chosen items (*β* = 0.26, *SE* = 0.10, *p* <.05), and decreased for correctly remembered rejected items (*β* = −0.17, *SE* = 0.08, *p* <.05) (Fig. 2) (all model results are reported in detail in Table S4 in, Supplemental Material).

**Figure 2.**
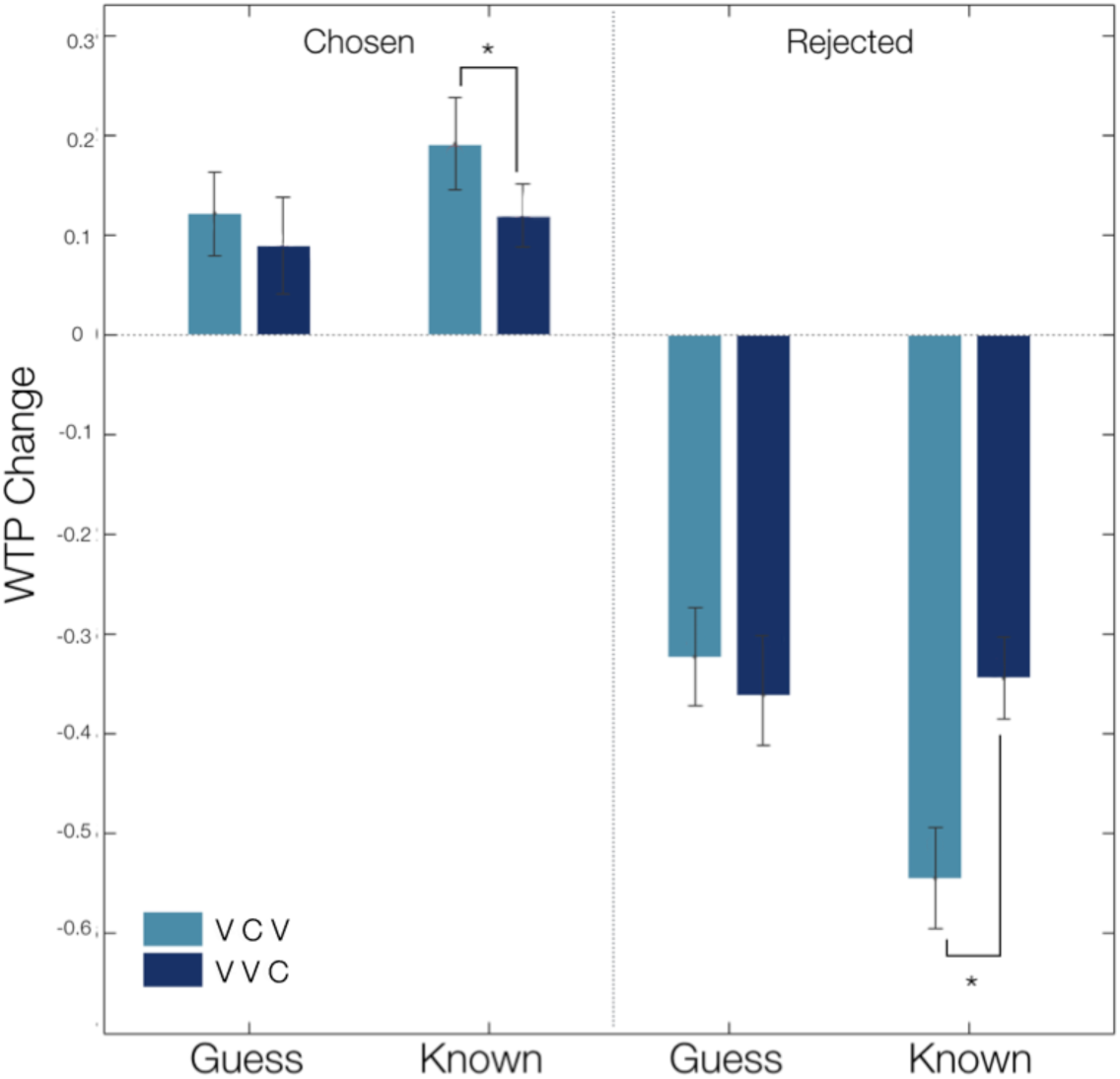
Behavioural results revealed memory-dependent, bidirectional choice-induced preference changes. For chosen/rejected remembered items, the change in willingness-to-pay (WTP) values was significantly higher/lower in the valuation-choice-valuation (VCV) condition as opposed to the valuation-valuation-choice (VVC) control condition. This effect did not reach significance for guessed choices.

### Neuroimaging Results for Control Analyses

#### Neural Representation of Monetary Value

We next verified that our WTP procedure engaged the ‘valuation network’, which relates to the encoding of subjective (monetary) value, i.e., preferences (Dolan and Dayan, 2013). To this end, we regressed participants’ trial-by-trial WTP scores against BOLD responses during the second valuation phase (GLM1, *Material and Methods*) from regions of interest (ROIs; Fig. S1) in vmPFC and vStr; Bartra, McGuire, and Kable 2013). We found a significant parametric modulation in both the vmPFC (MNI: 3 50 −4; *z* = 4.77; *p*_SV.FWE_ <.001) and vStr (MNI: 12 11 −1; *z* = 3.59; *p*_SV.FWE_ = .036) (Fig. S3).

#### Neural Correlates of Hard Choices

We then confirmed that hard choices activated decision-related brain regions to a greater extent than easy choices (GLM2, *Material and Methods*), suggesting decision conflict. A whole-brain analysis showed that activity in the left dorsal ACC [MNI: −6 26 4; extent threshold *p*_SV.FWE_ = .006 (*p*_uncorr._ <.001, height threshold); *z* = 4.21; *k* = 130] and left middle frontal gyrus [MFG, MNI: −45 23 26; extent threshold *p*_SV.FWE_ = .04 (*p*_uncorr._ <.001 height threshold); *z* = 3.91; *k* = 79] was significantly higher for hard compared to easy choices (Figure S3). These regions have previously been related to decisions between equally valued options, approach conflicts and choice anxiety (Kitayama et al., 2013; Shenav and Buckner, 2014; van Veen et al., 2009).

### Preference Changes following Hard Decisions

#### Neuroimaging Results

Next, we investigated the neural correlates of the memory-depended choice-induced preference changes at the time of re-evaluation. As a recent study (Chammat et al., 2017) provided initial evidence that this effect is associated with left hippocampus activity, we used their results to construct a ROI (Figure S1). Consistent with our behavioural results, the predicted critical interaction between experimental condition and remembered choice outcomes was associated with changes in left hippocampus activity (MNI: −24 −28 −16; *p*_SV.FWE_ = .012; *F* = 12.56, *z* = 3.18) (Figure 3D). An additional whole-brain analysis confirmed that no other brain regions showed this effect. Although the interaction between correctly guessed choice outcomes and experimental condition did not show any change in preference at the behavioural level, significant effects for correctly guessed items were found at the whole brain level in one cluster within left posterior parietal cortex (PPC), including the precuneus [MNI: −6 −55 32; extent threshold *p*_SV.FWE_ = .01 (*p*_uncorr._ <.001 height threshold); *z* = 4.61; *k* = 133] (Figure 3E), but not in the hippocampus.

**Figure 3.**
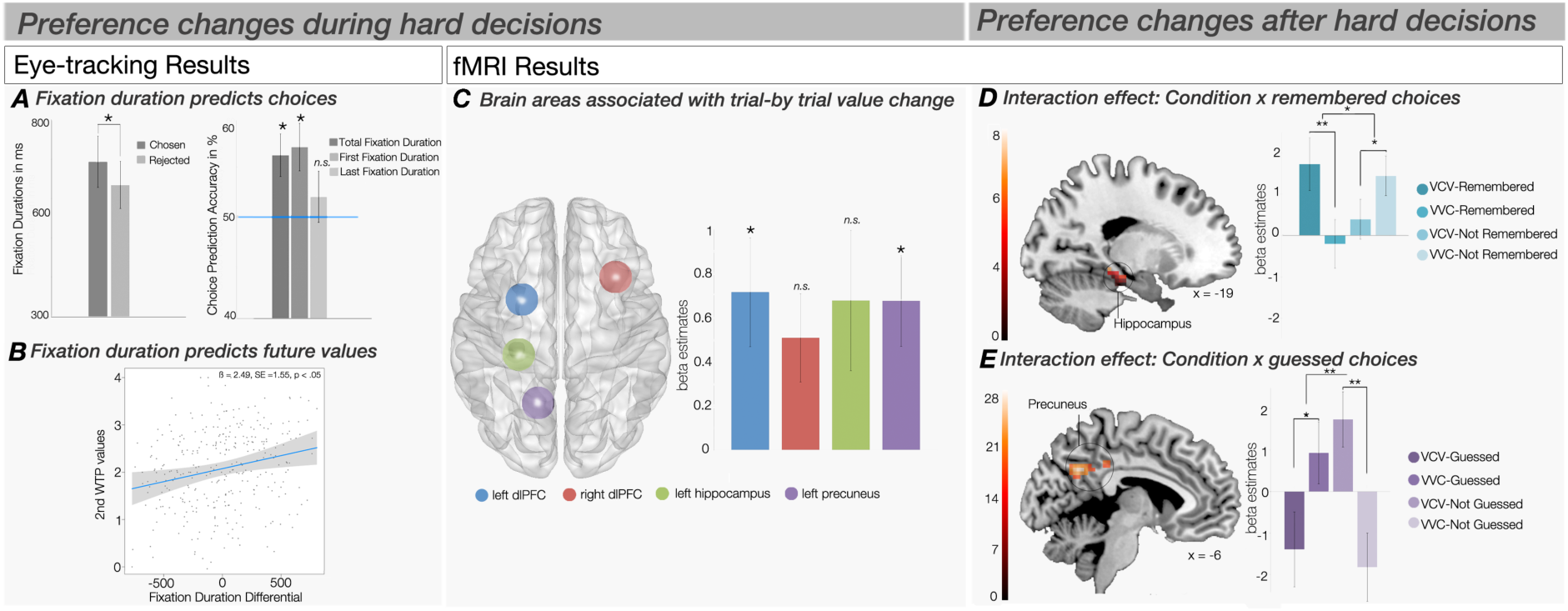
Preference formation during and following hard choices. In-choice results. (A) Fixation Duration for chosen and rejected items. (B) Choice prediction accuracy of total, first and last fixation duration. (B) Fixation duration differential (i.e., fixation duration of chosen items minus fixation duration of rejected item) predicted upcoming valuation change of chosen item. (C) Brain regions of interest associated with trial-by-trial preference change at the time of decision-making between equally valued items. Post-choice results. (D) The interaction effect between experimental condition and correctly remembered choice outcomes was associated with left hippocampus activity. (E) The interaction effect between experimental condition and correctly guessed choice outcomes was associated with left hippocampus activity.

### Preference Changes during Hard Decisions

#### Neuroimaging Results

Our crucial analyses involved testing whether preference changes were already present at a neural level *during* the process of making hard decisions, and whether these effects were moderated by memory processes, as suggested by our behavioural results. Previous studies suggested that the dlPFC (Izuma et al, 2010; Izuma et al., 2015), particularly the left (Harmon-Jones et al., 2015; Mengarelli et al., 2013), is directly involved in post-decisional preference changes. We therefore regressed trial-by-trial preference change scores for each of the hard choices as a parametric regressor against the BOLD data obtained during the first decision phase (GLM 3, *Material and Methods*) from predefined ROIs in the left and right dlFPC (Figure 3C and Figure S1) (Izuma et al., 2010). Our analysis showed that activity in the left dlPFC was predictive of subsequent preference changes for the later remembered items (MNI: −24 5 47; *p*_SV.FWE_ = .01, *z* = 3.61). The same contrast in right dlPFC ROI marginally missed the significance threshold (MNI: 33 20 30; *p*_SV.FWE_ = .07, *z* = 2.80).

As we found choice-induced preference changes to be encoded in hippocampus (remembered) and precuneus (guessed) for the second valuation phase, we further repeated the above analysis (GLM3) for the decision phase using these areas as ROIs (Figure S1). This analysis showed that the precuneus (MNI: −15 −58 29; *z* = 3.59; *p*_SV.FWE_ = .008), but not hippocampus (MNI: −27 −28 −19; *z* = 2.03; *p*_SV.FWE_ = .48) encoded the preference changes at the time of decision-making (Figure 3C).

To assess whether the effects were specific to the choice task stage in which choice can shape preferences (i.e., VCV), we repeated the analyses using the data obtained during second choice task stage (i.e., VVC). We did not find any significant results for any of the ROIs, indicating that the results were unique neural correlates of preference changes induced by choice and not artefacts due to regression to the mean (Chen and Risen, 2010).

#### Eyetracking Results

Our neuroimaging results do not address the question whether preferences changed as a consequence of choice, or alternatively, whether they changed during the decision-making *process*. To answer this question, we analysed fixation data during the decision making process (in the same phase 1) while participants were exposed to both options on the screen preceding their response.

*Fixation duration predicts choices.* The total fixation duration for chosen items (*M* = 693.59ms, *SD* = 253.34ms) was significantly higher than for rejected items (*M* = 634.65ms, *SD* = 232.80ms), (*t*(14) = 3.48, *p* = .004; *d* = 0.89). Similarly, the fixation duration ratio for chosen items (*M* = .44, *SD* = 0.15) was significantly higher than the fixation duration ratio for rejected items (*M* = .40, *SD* = 0.14, (*t*(14) = 2.89, *p* = .003; *d* = 1.06) (Figure 3A). A linear regression model (*Material and Methods*) showed that the probability of choosing an item was determined by its fixation duration ratio (*β* = 0.07, *SE* = 0.03, *p* <.05), but also by whether it was looked at first (*β* = 0.28, *SE* = 0.14, *p* <.05) and last (*β* = 0.37, *SE* = 0.14, *p* <.01). Both the accuracies with which choices could be predicted from the total fixation duration (*M* = .56, *SD* = 0.08) and the duration of the first fixation (*M* = .57, *SD* = 0.09) were significantly higher than chance (total fixation duration: *t*(14) = 2.91, *p* = .01; *d* = 0.75; first fixation duration: *t*(14) = 2.94, *p* = .01; *d* = 0.76). Fixation duration of the last fixation, however, did not predict choice outcomes (*M* = .52, *SD* = 0.1; *t*(14) = .72, *p* = .48) (Figure 3A).

*Fixation duration predicts updated preferences.* Next, we examined whether fixation durations also predicted subsequent changes in valuation for hard choices during the first decision phase. A LME model (*Material and Methods*) showed that the fixation duration differential (i.e., the mean difference between the fixation duration of chosen and rejected items) predicted the subsequent, updated WTP value for the chosen item (*β* = 5.55, *SE* = 1.96, *p* <.001, one-tailed), with higher fixation rates being linked to higher updated values, as early as 1000ms after stimulus presentation. This finding also held when controlling for the initial WTP values (*β* = 2.49, *SE* = 1.55, *p* <.05, one-tailed) (Figure 3B). The analysis revealed no interaction effect with item memory.

## Discussion

Preference changes following hard choices have been interpreted as a result of cognitive dissonance reduction after decisions have been made and preferences are re-assessed (Festinger, 1957). As such, previous fMRI studies of preference changes induced by hard choices solely focused on neural correlates of preference changes during the re-evaluation of alternatives (Chammat et al., 2017; Izuma et al., 2010). Our study is the first to reveal that preference changes were linked with neural activity much earlier, i.e., *during* hard decisions are being made. Specifically, activation in a brain network comprising left dlPFC and precuneus was predictive of upcoming preference changes effects. Further, we found that fixation durations predicted choices as well as future valuations. These outcomes point to a profoundly different mechanism of preference change in which preferences are adjusted online while decision-makers are deciding among equally valued alternatives.

Our behavioural results confirmed that for such equally valued, consumable items, the valuations increased after choosing and decreased after rejecting when valuations were measured by incentive-compatible willingness-to-pay assessments (Voigt et al., 2017). Consistent with other earlier reports (Chammat et al., 2017; Salti et al., 2014) we refined these results by showing that preferences changed only for choices that were explicitly remembered later. However, these earlier studies could not rule out the possibility that participants did not actually remember which choices they made earlier, but simply inferred (i.e., guessed) them based on their updated preferences. By asking participants to label their choice outcomes as ‘remembered’ or ‘guessed’ we could show that explicit choice memory, but not correct guessing, was linked to behavioural choice-induced preference change effects.

We further replicated Chammat and colleagues’ (2017) findings that memory-depended choice-induced preference changes were associated with left hippocampus activity – a core region involved in long-term episodic memory (Bird and Burges, 2008). In our study, the same neural correlates of the spread of alternatives was only found for remembered items. In addition, while preference changes were absent at the behavioural level for correct guesses, a significant neural effect for spread of alternatives in this condition could nevertheless be observed in the precuneus. This region has been associated with the rapid formation and retrieval of episodic memory (Brodt et al., 2016), with self-relevant processing (Kircher et al., 2002) and with decisions based on guessing (Bode, Bogler, and Haynes, 2013). The precuneus might therefore be involved in less certain retrieval processes for items, which only lead to smaller, behaviourally sub-threshold choice-induced preference change effects.

Previous explanations for the observed spread of alternatives following hard choices (Chammat et al., 2017; Izuma et al., 2010; Salti et al., 2014) were in accord with prominent theories that valuations are adjusted post-hoc to match previous choices (e.g., cognitive dissonance theory, Festinger, 1957; self-perception theory, Bem, 1967). However, the temporal dynamics of choice-induced preference changes were never explicitly tested. Some other studies attempted to investigate whether preference change during the initial decision phase (Colosio et al., 2017; Jarcho et al., 2011; Kitayama et al., 2013) but remained inconclusive as they did not distinguish decision conflict from preference change (Colosio et al., 2017) did not control for potential regression to the mean artefacts (Chen and Risen, 2010) or used noisy, incentive-incompatible preference assessments, which might be ill-suited to investigate choice-induced preference changes (Voigt et al., 2017). Our study accounted for these methodological issues, and clearly showed that trial-by-trial preference changes were already reflected in the dlPFC during the decision process. The left dlPFC has been shown previously to be involved in the implementation of preference change after hard choices were made (Izuma et al., 2010; Mengarelli et al., 2013). Here, we extended these findings in showing that this area was involved much earlier. In addition, we tested whether the memory-related regions, which were found to reflect the spread of alternatives in our study in the re-valuation phase (as conceptualised by Chammat et al., 2017) also tracked changes in preferences during decision-making. Interestingly, such effects were absent in the hippocampi but present in the precuneus.

Taken together, our fMRI results suggest a process in which first, a decision conflict among equally valued alternatives is detected, based on subjective values of the choice alternatives. In line with this, monetary values were associated with activity in vmPFC and vStr in our study (cf., Bartra et al., 2013) while decision conflict was reflected in enhanced activity in the ACC and MFG (cf., Botvinick, 2007; Shenav and Buckner, 2014). The detection of decision conflict could then trigger an updating process for stimulus values during decision formation, possibly to resolve the initial conflict and to avoid similar near-stalemate situations in the future. This idea is consistent with a revised version of cognitive dissonance theory (Harmon-Jones et al., 2015) which states that a decision conflict needs to be resolved first in order to enable the individual to prepare a choice plan. This process could therefore involve the dlPFC, which is strongly related to decision-making and working memory (Yan, Wei, Zhang, Jin, and Li, 2016) as well as the precuneus, which might be more involved in the initial formation of episodic memory and potentially driving self-referential decision processes via allocation of attention (Brodt et al., 2016; Kircher et al., 2002). Shifts in spatial attention related to precuneus activity during decision formation could then feed into the reconstruction of new value information in the dlPFC, which in turn could store the new value representation in working memory, assisting the optimal decision between the options. This is in line with demonstrations that value reconstruction evolves from posterior parietal to dorsolateral prefrontal regions (Harris, Adolphs, Camerer, and Rangel, 2011). Consequently, during subsequent re-valuation, stronger changes in preference, and also stronger memory-related signals, would be found for the same items, which is what we and others (Chammat et al., 2017; Salti et al., 2014) observed.

In order to investigate whether preferences were indeed reconstructed during decision formation for hard decisions, we also analysed fixations via eye-tracking. Previous studies postulated a causal link between visual fixations and the formation of subjective values and value-based choice (Krajbich et al., 2010; Shimojo et al., 2003). According to these studies, the allocation of spatial attention might lead to an increase in information accumulation in favour for the fixated object and, in turn, a higher likelihood of this object to be chosen. In accordance with these reports, we found that fixation duration of the first fixation and total fixation duration predicted choices. Beyond choice itself, fixations also predicted changes in future values for chosen items. This analysis could not explicitly take into account regression to the mean effects, but given that we demonstrated clear choice-induced preference change effects after controlling for such effects for the same items, this strongly suggests that our findings did indeed reflect preference adjustments during hard decision. As fixation durations provide a window into this decision process, these findings suggest that during decision formation, the future state of preference representations were dynamically reconfigured. These reconfiguration processes potentially also drive the encoding of choice options in episodic memory, meaning that subsequent choice memory might not reflect random variations in retrieval strength, but systematic differences in encoding strength during decision formation.

### The Formation of Decisions and Preferences

Our results cannot unambiguously disentangle which processes contribute to making the decision versus adjusting preferences. As early fixations predicted choices, it is possible that these fixations reflect allocation of attention, potentially acting as a ‘symmetry breaker’ when confronted with equally valuable options. This could be driven by the precuneus, which has been related to choices under indifference (Bode et al., 2013; Soon, He, Bode, and Haynes, 2013). The precuneus also has major subcortical connections to the pretectal area and the superior colliculus (Leichnetz, 2001; Yeterian and Pandya, 1993), which contribute to attentional shifts via eye movement control (Moschovakis, 1996). However, early fixations patterns must not necessarily be the result of random processes, but could themselves be driven by exogenous stimulus properties, such as its saliency (Itti and Koch, 2001), or residual differences in subjective value, which were not adequately captured by our WTP measurements. Further, it is possible that for the longer fixated item, more choice-attributes were considered, leading to a choice-advantage and preference increase (Orquin, Mueller, and Loose, 2013). Alternatively, the mere exposure effect (Zajonc, 1968), which states that preferences increase as a function of exposure duration, might also be relevant here.

In conclusion, our findings support a dynamic view of preference formation during decision-making, enabling the individual to make a value-based choice. Future studies are needed to explore what factors underlie the early fixations, which predicted both choice and updated preferences, and whether similar effects can be found beyond value-based choice.

## Materials and Methods

### Participants

The analyses were based on data from 22 participants (13 females; age 18 to 37 years; *M* = 23.57, *SD* = 4.93). Participants were right-handed English speakers with (corrected-to-) normal vision, who fasted for four hours prior to the study. Eye-tracking data was acquired for 15 participants (full overview provided in Table S1 in Supplementary Material and Methods). The study was approved by the University of Melbourne Human Research Ethics Committee (no. 1442440).

### Experimental Task and Procedures

The experiment consisted of two main consecutive tasks: *the incentive-compatible free choice task*) (Voigt et al., 2017) followed by a *choice memory task*. The first task consisted of four task phases: valuation phase 1, decision phase 1, valuation phase 2, and decision phase 2 (Fig. 1). Neuroimaging data was acquired during the decision phases and the second valuation phase. Eye-tracking data were acquired during the first decision phase.

### The incentivised free-choice Task

*(i) Valuation phase 1:* Each trial (292 total trials) started with a central fixation cross (jittered between 1s and 3s), which was followed by a pseudo-randomly selected snack food stimulus (1s) (Supplementary Material and Methods). Subsequently, participants indicated how much they were willing to pay for that item on a continuum from $0-$4. Responses were measured by moving a graphical slider along a continuous valuation scale. All responses were made via a MRI-compatible fibre optic trackball and were restricted to 3s.

*(ii) Decision phase 1*: A maximum of 80 ‘hard’ and 40 ‘easy’ choice pairs were created, based on the responses of the valuation phase 1, by pairing either items with highly similar (hard) valuations or dissimilar (easy) valuations, respectively (Supplementary Material and Methods). Half of the hard and easy choice pairs (60 total trials) were shown in a pseudo-randomised order, requiring participants to make binary (two-alternative forced-choice; 2AFC) decisions for the item they preferred (Table S3 in Supplementary Material and Methods). Trials were presented into two separate runs (i.e., 30 trials per run) to allow for a short break. Each trial started with a short fixation period (jittered between 4 and 9s), followed by a snack food pair. Critically, it was emphasised that only the item they chose could feature in a subsequent Becker-DeGroot Marshak (BDM auction). In the BDM auction (Supplementary Material and Methods) participants had the chance of buying one of the chosen items, and their WTP would serve as the bids from their pre-allocated budget of $4. As such, bids and decisions were incentivised and consequential (Voigt et al., 2017). The response window was 3s after which the selected choice was highlighted by a black frame for 1s.

*(iii) Valuation phase 2:* This task phase was identical to the valuation phase 1.Participants were instructed that the purpose was not to probe their memory of the first valuation, but to provide another, independent valuation.

*(iv)Decision phase 2:* This task phase was identical to the first decision phase with the only difference that the remaining, unused 60 choice pairs were presented (Table S3, Supplementary Material and Methods). This allowed us to use the first and second valuations for these items as a control sequence (VVC), assessing changes in valuation that were attributable to regression-to-the-mean effects (Chen & Risen, 2010).

#### The choice memory task

This task was completed outside the MRI scanner, and in 240 trials, participants were sequentially presented with all snack foods again. They indicated whether they remembered having previously chosen or rejected it. Critically, participants were asked to distinguish whether they were absolutely certain (options: *Chose!* or *Rejected!*; trials labeled “remembered”), or whether they felt that they were guessing (*Chose?* and *Rejected?;* trials labeled “guessed”) their response. In addition, participants could also indicate that they believed that the item did not feature in the experiment (i.e., *No option!* or \ *No option?*). If participants selected these options, these trials were coded as *falsely remembered* or *falsely guessed*. Participants were unaware of the subsequent memory task throughout the fMRI experiment; that is, they were not explicitly instructed to memorise their choices.

### Functional MRI Data Acquisition and Analysis

Functional MRI data acquisition and pre-processing were carried out using standard procedures described in (Supplementary Material and Methods). In order to address our research questions, five general linear models (GLMs) were constructed at the participant level (a detailed description can be found in (Supplementary Material and Methods). GLM 1 modelled the *neural representation of WTP value* for the second valuation stage. This contained two regressors of interests: (i) an onset regressor for the presentation of food stimuli (ii) and a parametric regressor for participants’ WTP response (ranging from $0 to $4) for that food stimulus. GLM 2 modelled the *neural correlates of hard decisions* and established where in the brain choice difficulty was represented during the crucial first decision phase. The model included two main regressors of interest: (i) onset for hard decisions and (ii) onset of easy decisions. GLM 3 modelled *memory-dependent choice-induced preference changes* effects in second valuation phase, closely following the approach by Chammat et al. (2017): trials were sorted into four conditions: (i) remembered items in the VCV condition, (ii) forgotten items in the VCV condition, (iii) remembered items in the VVC condition, (iv) forgotten items in the VCV condition. We regressed the main effect of condition, the main effect of correctly remembered choice outcomes and the interaction between condition and correctly remembered choice outcomes using a flexible factorial model in bilateral hippocampus. GLM 4 explored the possibility of preference changes for correctly guessed trials. We constructed a similar model to GLM 3, but with guessed trials only. GLM 5 was used to model *choice-induced preference change effects during hard choices* in the *a priori* defined ROI in the dlPFC. For this GLM, we went beyond the model suggested by Chammat et al. (2017) that only considered the spread of alternatives but entered the PCS scores as a parametric predictor for hard decision items into the model. For all GLMs, regressors were convolved with a canonical hemodynamic response (HRF) and together with the motion parameters from the realignment procedure regressed against the BOLD signal in each voxel. Results for ROIs were assessed via small volume correction (SVC). For whole-brain analyses correction for multiple comparisons [family-wise error (FWE), *p* <.05] was performed at cluster level.

### Eye-tracking Acquisition and Analysis

Right eye movements were recorded for the first choice task stage via an MR-compatible infrared video-based system measuring corneal reflection (Eyelink 1000) at 500Hz. Analyses were conducted in Matlab and R. To assess whether fixation parameters predicted choice, a GLM was conducted. The fixation duration ratio, first and last fixation count of the chosen item were regressed on choice outcome. To determine whether fixation duration predicted updated preferences, we conducted another GLM and regressed the fixation differential (i.e., the mean difference between the fixation duration of chosen and rejected items) and the initial WTP values of the chosen item on the chosen item’s WTP values from the second evaluation phase. For this GLM, we analysed the data from 1000ms post-stimulus presentation, as previous research reported value-formation at this later stage (Harris et al., 2011)

## Acknowledgements

The authors thank Simon Lilburn, and Jacob Paul for helpful discussions and Sophia Bock, William Turner and Richard McIntyre for support with MRI data acquisition. This study was supported by an Australian Research Council Discovery Early Career Researcher Award (DE 140100350) to S.B.

## Notes

**Declaration of Interests:** The authors declare no declarations of interest.

